# The unique pattern of mannose-capped lipoarabinomannan expression in *Mycobacterium tuberculosis* with different drug resistant profiles following isoniazid stress

**DOI:** 10.1101/2021.10.20.465078

**Authors:** Manita Yimcharoen, Sukanya Saikaew, Usanee Anukool, Ponrut Phunpae, Sorasak Intorasoot, Watchara Kasinrerk, Chatchai Tayapiwatana, Bordin Butr-indr

**Author notes:** Corresponding author, (B. Butr-Indr).

## Abstract

Tuberculosis (TB) is a global health problem caused by *Mycobacterium tuberculosis* (MTB) infection. The main problem of TB treatment is the emergence of drug resistance, which can occur by inappropriate of antibiotic used. Isoniazid (INH) is the first-line anti-TB drug that inhibits mycolic acid synthesis, an important part of the mycobacterial cell wall. Mannose-capped lipoarabinomannan (ManLAM) is an essential cell wall part that plays a role as an immunomodulator and acts as a virulence factor. In this study, MTB clinical isolates with different drug resistant profiles were used to determine the expression of ManLAM related genes including *pimB, mptA, mptC, dprE1, dprE2 and embC* by qRT-PCR. Stress-related genes including *hspX, tgs1*, and *sigE* were determined by multiplex real-time PCR with probe assay. Sanger sequencing of ManLAM related genes and genes associated with drug resistance (*inhA, katG*, and *rpoB*) were analyzed. In response to INH, the expression pattern of ManLAM related genes was different among four strains. Interestingly, MDR-TB markedly up-regulated ManLAM related genes greater than others. Stress-related genes *hspX* and *tgs1* were significantly upregulated in MDR response to INH, whereas *sigE* was significantly upregulated in MDR response to RIF and INH-R. DprE1 is crucial for MTB and it is a valuable target for anti-TB drugs. RIF-R and MDR isolates show C→T mutation at nucleotide position 459 of the *dprE1* gene leading to the same amino acid at codon 153. Codon usage analysis for DprE1 showed that RIF-R and MDR preferred ACT codon over drug sensitive strains. This work provides the expression pattern of ManLAM related genes and stress responder genes, which are key factors in the interaction between MTB and host. Moreover, ManLAM is a possible factor that plays an important role in the adaptive mechanism and the drug resistance mechanism of mycobacteria.

**Author summary:** The adaptive mechanism of mycobacteria in response to stressors is an important strategy to promote their virulence and pathogenesis. This study determined the effect of antibiotic stress on *Mycobacterium tuberculosis* (MTB) focusing on mannose-capped lipoarabinomannan (ManLAM), which is one of the virulence factors that modulate host immune response. Multiplex real-time PCR with probe assay targeting stress responder genes and qRT-PCR targeting ManLAM related genes were performed. Isoniazid acts as a stressor to induce stress response in mycobacteria, as shown in the up-regulation of stress-related genes including *hspX, tgs1*, and *sigE*. The expression pattern of ManLAM related genes in drug resistant and drug sensitive-MTB in response to INH was different, causing a unique pattern. ManLAM related genes respond to isoniazid mostly in drug resistant strains and are present at high expression levels in INH-R and MDR. The results suggest that ManLAM is one factor involved in the adaptive mechanism of MTB response to antibiotic stress and probably associated with the emergence of MTB drug resistance. This work provides new insights into the adaptive mechanism of mycobacterial response to isoniazid that will improve understanding of how mycobacteria develop drug resistance.

## Introduction

*Mycobacterium tuberculosis* (MTB) is an infectious agent of tuberculosis (TB), which continues to be a global health problem due to the emergence of drug-resistant strains. There are several factors that support the growth of MTB and TB pathogenesis. The cell wall of mycobacteria plays a protective role against toxic compounds or antibiotic agents [1]. Mannose-capped lipoarabinomannan (ManLAM) is an important glycolipid of the MTB cell wall, which is a virulence factor [2]. ManLAM has an immunomodulatory effect on the host immune response in a variety of ways, including cytokine production, phagosome maturation, antigen presentation, T cell activation, and polarization [3]. The structure of ManLAM consists of a mannosyl-phosphatidyl-myo-inositol (MPI) anchor linked to a mannan backbone, an arabinan domain and mannose caps motifs [4]. The biosynthesis of ManLAM follows a pathway from phosphatidyl inositol (PI), phosphatidylinositol mannosides (PIMs), lipoarabinomannan (LM), lipoarabinomannan (LAM) and mannose-capped lipoarabinomannan (ManLAM), respectively [5]. Several genes are responsible for the biosynthesis of ManLAM, including *pimB, mptA, mptC, dprE1, dprE2*, and *embC*. pimB (Rv2188c) is responsible for the synthesis of PIMs involved in the early step of ManLAM biosynthesis [6]. MptA (Rv2174) is defined as α-(1→6)-mannosyltransferase responsible for the elongation of mannan backbone of LM [7]. MptC (Rv2181) is defined as α-(1→2)-mannosyltransferase responsible for the branching of LM and required for mannose capping [8]. DprE1 (Rv3790) and DprE2 (Rv3791) are responsible for the epimerization of decaprenylphosphoryl ribose (DPR) to decaprenylphosporyl arabinose (DPA), a precursor for LAM [9]. embC (Rv3793) has been identified as an arabinosyltransferase involved in the biosynthesis of branched arabinan polymers on arabinogalactan and LAM using DPA as a precursor [10]. The adaptive ability to respond to the stressful host environment is a critical strategy for MTB to evade host killing mechanisms and promote disease progression [11, 12]. They regulate the expression of virulence genes, modulate cell wall components and change their antibiotic susceptibility profile in response to host-derived stresses [12-15]. hspX (*acr*, Rv2031c) known as alpha-crystallin is crucial for MTB survival in the host stress environment and during the dormancy phase [16-18]. Previous studies have shown that *hspX* expression is induced by low oxygen tension or hypoxia [18, 19], high nitric oxide and carbon monoxide concentrations [18], and a multiple stress model [12] [low oxygen (5%), higher levels of carbon dioxide (10%), low nutrient (10% Dubos medium), and acidic pH (5.0)]. tgs1 (Rv3130c) is defined as a triacylglycerol synthase involved in the synthesis of triacylglycerol (TG), which is required for long-term survival [20]. High expression of *tgs1* and accumulation of TG have been found under stress conditions [20] and in a multiple stress model [12], resulting in drug-tolerant MTB. sigE (Rv1221) is an alternative RNA polymerase sigma factor that is regulated in response to several stress such as hypoxia [19, 21], nutrient starvation [22] and acidic pH [23, 24]. These data suggest that *hspX, tgs1* and *sigE* are stress-related genes and can be used as stress markers. Adaptive mechanisms of MTB associated with acquired antibiotic resistance may be due to inappropriateness of the antibiotic used [25]. Isoniazid (INH) is a first-line drug that requires activation by redox enzymes encoded by *katG* gene of MTB to inhibit enoyl-acyl carrier protein reductase (InhA) resulting in inhibition of mycolic acid synthesis [26], an essential component of cell wall. INH treatment is one type of stress inducer (antibiotic stress) that may alter other parts of MTB in addition to mycolic acid synthesis. A microarray hybridization study of INH treatment showed that INH-treated MTB upregulate several genes encoding proteins involved in fatty acid biosynthesis, trehalose dimycolyl transfer, fatty acid degradation, peroxidase activity, transport, and efflux pump [27]. Analysis of total lipids of MTB clinical isolates showed that clinical isolates of INH-resistant strains had significantly lower levels of phthiocerol dimycocerosate (PDIM), trehalose monomycolate (TMM), and phosphatidylinositol mannosides (PIMs) compared with INH-sensitive strains [28]. INH treatment causes free trehalose accumulation, which is a signal for *IniBAC* regulator (IniR) to induce the major cell wall stress operon *iniBAC* [29]. These data suggest that INH treatment affects the expression of genes involved in the biosynthesis of mycobacterial cell wall components.

In this study, MTB with different drug resistance profiles including drug-sensitive (H37Rv), mono-isoniazid resistance (INH-R), mono-rifampin resistance (RIF-R) and multi-drug resistance (MDR), were treated with or without INH C_Max_ (6 µg/ml) [30] concentration for 30 min to determine the effect of INH on ManLAM related genes expression. The relative expression of ManLAM related genes such as *pimB, mptA, mptC, dprE1, dprE2*, and *embC* was determined by qRT-PCR. To determine the relationship between genetic background and phenotypic phenomenon, a genetic analysis of ManLAM related genes and drug resistance associated genes was performed. In addition, the expression of stress-related genes such as *hspX, tgs1* and *sigE* induced by antibiotic stress INH or RIF was investigated in drug-sensitive and drug-resistant strains (INH-R and MDR) to study the effects of antibiotic stress on MTB using stress-related gene markers.

## Materials and methods

### Bacterial strain and cultivation conditions

All mycobacteria used in this work, including H37Rv (Drug sensitive strain), INH-R, RIF-R, and MDR, were obtained from the Office of Disease Prevention and Control, 1 (Chiang Mai, Thailand). The drug susceptibility profile was estimated and reported in a previous publication [31]. Mycobacteria isolates were cultured in Lowenstein-Jensen medium (LJ medium) (Biomedia, Thailand) at 35 °C for 4 weeks.

### Drug susceptibility test (DST)

To confirm the phenotypic drug resistance profile of MTB isolates, the agar proportion method was performed according to the Clinical and Laboratory Standards Institute (CLSI) document M24-A2 [32] using recommended critical concentrations of isoniazid (0.2 and 1 μg/ml), rifampin (1 μg/ml), streptomycin (2 μg/ml), and ethambutol (5 μg/ml).

### DNA extraction

MTB colonies were resuspended in 25 µl of 1 mg/ml lysozyme and incubated in water bath at 37°C for 10 min. Twenty-five microliters of proteinase K and 75 µl of 0.1 M Tris HCl were added and incubated in water bath at 37°C for 10 min. Then it was incubated in a hot block at 95°C for 10 min. The amount of DNA was determined by measuring the optical density of 260 and 280 nm using a UV Spectrophotometer (Biotech Epoch™) and kept at 4 or-20 °C before use.

### GenoType MTBDRplus assay

To test the genotypic drug resistance profile of MTB isolates, the GenoType MTBDRplus assay (Hain Lifescience, Germany) was performed according to the manufacturer’s recommendations.

### Antibiotic preparation

Isoniazid (INH) (Sigma-Aldrich, USA) and rifampin (RIF) (Applichem, Germany) were prepared according to a previous publication [31]. INH and RIF were prepared at a stock concentration of 1 mg/ml, sterilized with 0.2 µm nylon membrane filters, and dissolved in sterile distilled water. Stock solutions were stored at −20 °C prior to use.

### Antibiotic treatment conditions

Each mycobacterium was cultured and treated following previous publication [31]. Mycobacteria were culture in 20 ml of Middlebrook 7H9 (M7H9) supplement with 0.05% (v/v) tween 20, 10% (v/v) OADC and 0.5% (v/v) glycerol at 35 °C for 6 days. Prior drug treatment, 10 ml of bacterial cultures were transferred to new tube and adjusted to McFarland standard No.1 with Middlebrook 7H9. Isoniazid at a C_Max_ concentration (6 µg/ml) or rifampin at a C_Max_ concentration (24 µg/mL) [30] was added to each mycobacterial culture tube, mixed and incubated at 35°C for 30 min. Untreated tube of each strain was used as control. The treated cells were pelleted by centrifugation at 12,000×g at 4 °C for 20 min (Beckman Allegra X-15R) and the supernatant was discarded. The pellets were washed twice with sterile phosphate buffered solution (PBS) and centrifuged at 12,000×g at 4 °C for 15 min before performing RNA extraction.

### RNA extraction and complementary DNA (cDNA) conversion

RNA extraction was performed using Nucleospin^®^ RNA kit (MACHEREY-NAGEL, Germany) following the manufacturer’s guidelines. Mycobacterial cells were resuspended in 100 μl of Tris-EDTA buffer (10mM Tris-HCl, 1mM EDTA; pH 8) containing 2 mg/ml of lysozyme for 5 min. Cells were lysed with 350 μl of Buffer RA1 provided in Nucleospin^®^ RNA kit and 10mM of dithiothreitol, mixed by repetitive pipetting. The suspension was transferred to a sterile screw cap microcentrifuge tube containing 0.1 mm-size zirconia-silica bead (BioSpec, Oklahoma). Mycobacterium cells were disrupted three times by OMNI Bead Ruptor (OMNI, USA) at speed No. 2 for 1 min and incubated at room temperature (RT) for 5 min. After the incubation step, RNA isolation was further performed by recommended protocols (MACHEREY-NAGEL, Germany). cDNA conversion was performed from each sample by using ReverTra Ace^®^ qPCR RT Master Mix with gDNA remover (Toyobo, Japan). The amount of RNA and DNA was determined by measuring the optical density of 260 and 280 nm using a UV Spectrophotometer (Biotech Epoch™) and samples were kept at 4 or-20 °C before use.

### Polymerase Chain Reaction (PCR) and DNA Sequencing analysis

To study genetic variation of LAM related genes (*pimB, mptA, mptC, dprE1, dprE2*, and *embC*) and genes associated with drug resistance (*inhA, katG*, and *rpoB*). Fifty ng/µl of genomic DNA was used as a DNA template. PCR was performed by using KOD One™ PCR Master Mix-Blue (Toyobo, Japan), final volume of 50 µl following the manufacturer’s instructions, and the Lab Cycler Basic-Thermal Cycler (SensoQuest, Germany) was used as a PCR machine. Primer sequences are shown in Table 1 and PCR conditions are shown in Table 2. After the PCR process, agarose gel electrophoresis was performed to investigate PCR product size. The single band as their expected product size without non-specific bands demonstrates the specificity of designed primers and appropriate PCR conditions. The NucleoSpin^®^ Gel and PCR Clean-up (MACHEREY-NAGEL, Germany) were used to purify the PCR products. Next, PCR products were subjected to DNA sequencing by the automated sequencer ABI Prism 3730 XL, using the same primers as used for amplification. Sequence alignment was performed using ClustalW in the BioEdit program. *Mycobacterium tuberculosis* H37Rv (GenBank: CP003248.2) was used as a reference strain. Codon usage analysis was performed by using visual gene developer 1.9 (Build 785) program.

**Table 1.**
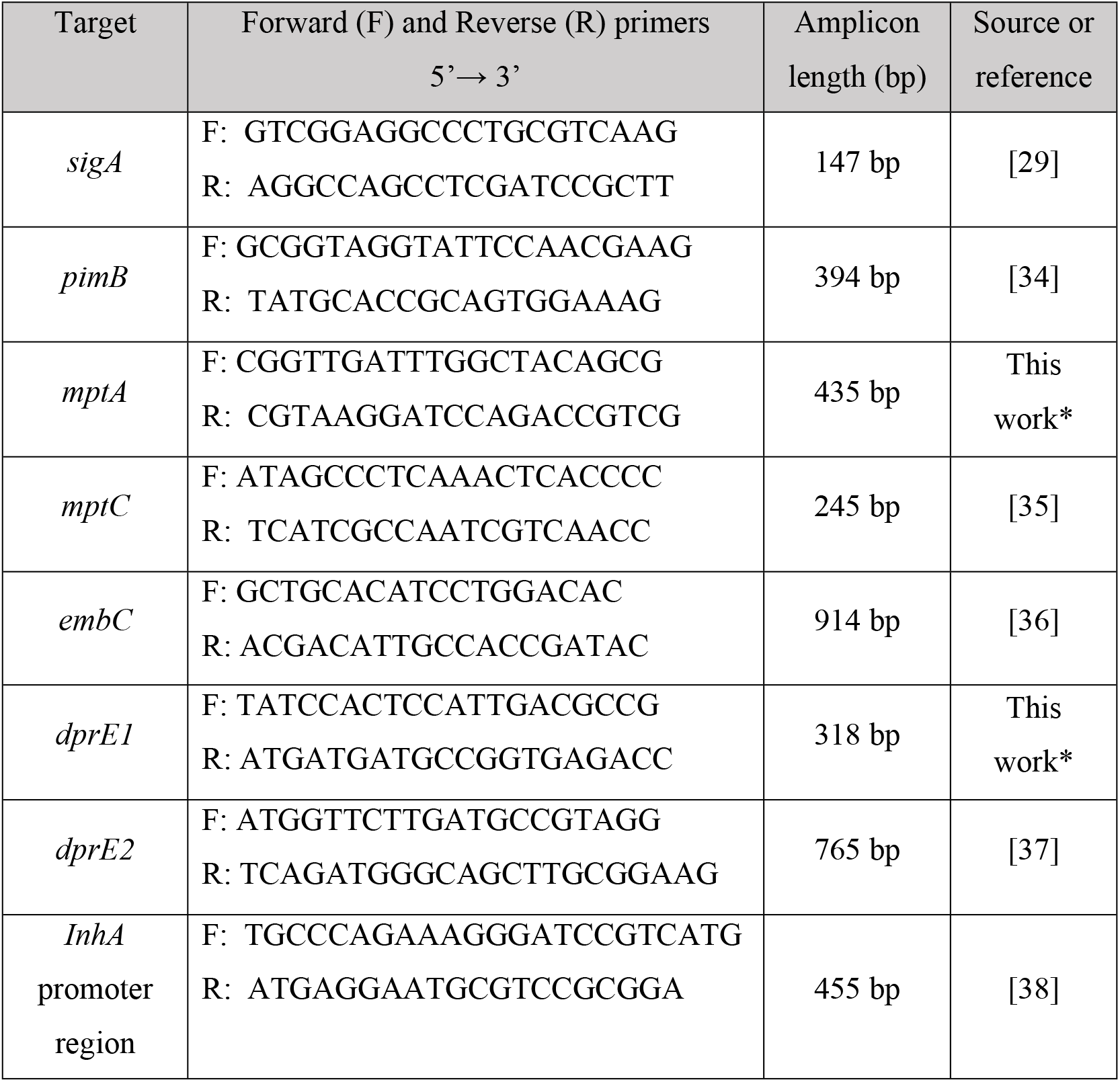
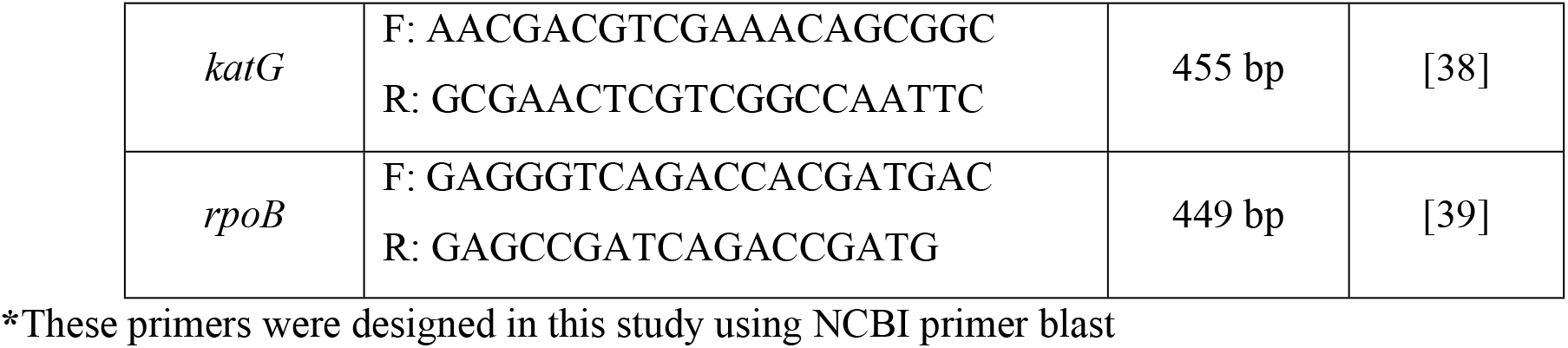
Primer sequences for ManLAM related genes and genes associated with drug resistance.

**Table 2.** PCR conditions for DNA sequencing.

| Steps | Temperature (°C) | Time | Cycles |
| --- | --- | --- | --- |
| Pre-denaturation | 95 | 5 min | 1 |
| Denaturation | 95 | 30 sec | 35 |
| Annealing | 68 (*inhA* and *katG*), 65 (LAM related genes), 56 (*rpoB*) | 15 sec | 35 |
| Extension | 72 | 10 sec | 35 |
| Final extension | 72 | 5 min | 1 |

### qRT-PCR analysis target LAM related genes

To quantify the expression of LAM related genes of MTB induced by INH treatment. The cDNA of treated-MTB and untreated-MTB were used as a template. The specificity test of each primer was confirmed by PCR and agarose gel electrophoresis. Real-time PCR was performed by using THUNDERBIRD^®^ SYBR^®^ qPCR Mix (Toyobo, Japan) in the CFX96 Touch Real-Time PCR Detection System (Bio-Rad, LABORATOIRES INC.). Primer sequences are shown in Table 1, and PCR conditions are shown in Table 3. The fold change of gene expression was calculated by the 2^−ΔΔCT^ method [33]. The house keeping gene, *sigA* (Rv2703c) [31] was used to normalize expression levels of each interested gene.

**Table 3.** PCR conditions for ManLAM related genes expression analysis.

| Steps | Temperature (°C) | Time | Cycles |
| --- | --- | --- | --- |
| Pre-denaturation | 95 | 5 min | 1 |
| Denaturation | 95 | 30 sec | 40 |
| Annealing | 60 | 30 sec | 40 |
| Extension | 72 | 30 sec | 40 |
| Final extension | 72 | 5 min | 1 |

### Multiplex real-time PCR with probe assay target stress-related genes

To quantify the expression of stress-related genes of MTB induced by anti-TB drug, isoniazid or rifampin, multiplex real-time PCR with probes was designed and used in this work. Primers and probes were designed using NCBI blast and PrimerQuest tool except *hspX*. Primers and probes were listed in Table 4 and Table 5, respectively. PCR conditions were shown in Table 6. The fold change of gene expression was calculated by the 2^−ΔΔCT^ method [33]. The house keeping gene, *sigA* (Rv2703c) [31] was used to normalize expression levels of each interested gene. The PCR product was run on electrophoresis to confirm size and specificity of the designed primers.

**Table 4.**
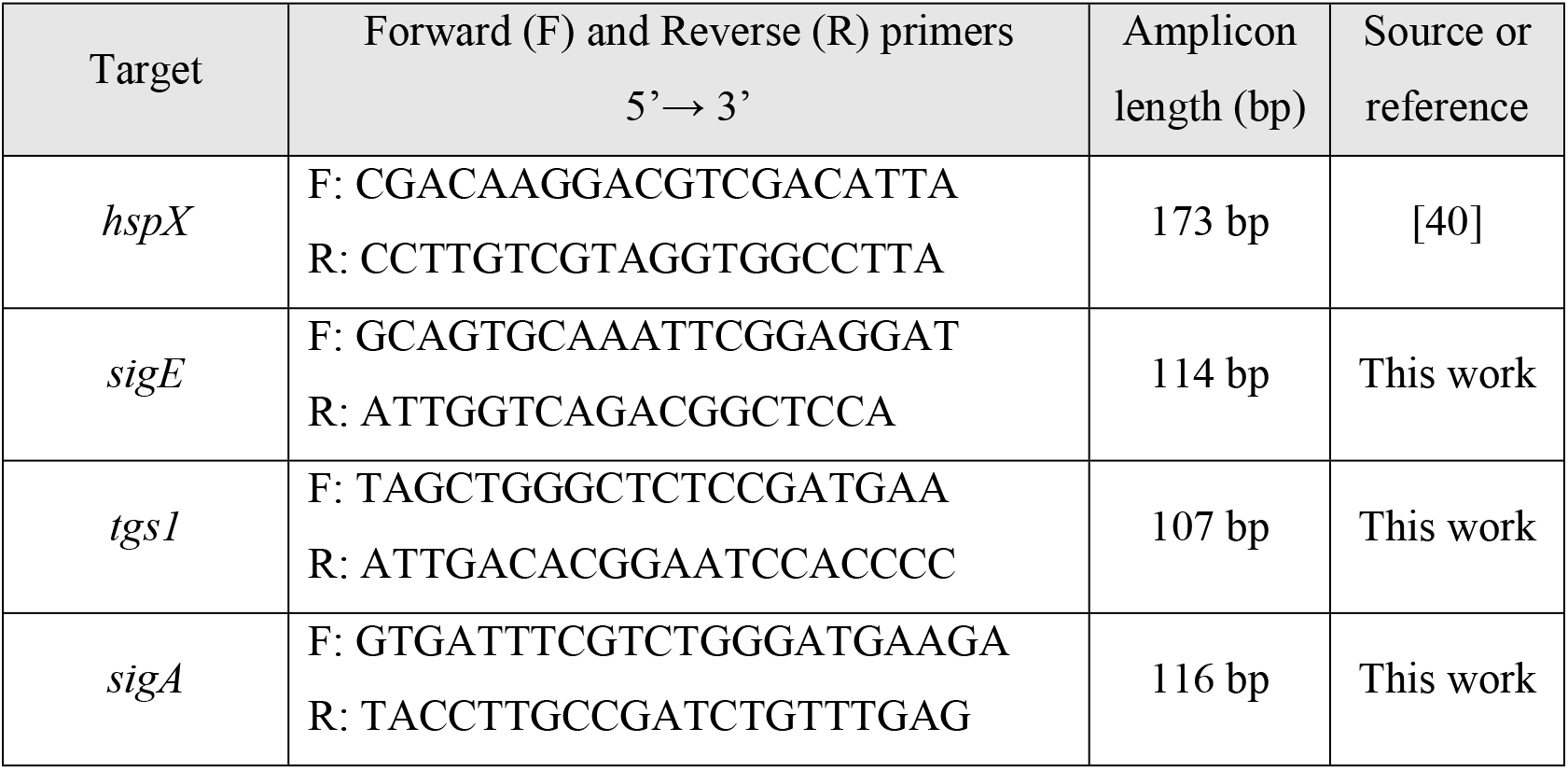
Primer sequences for multiplex real-time PCR target stress-related genes.

**Table 5.**
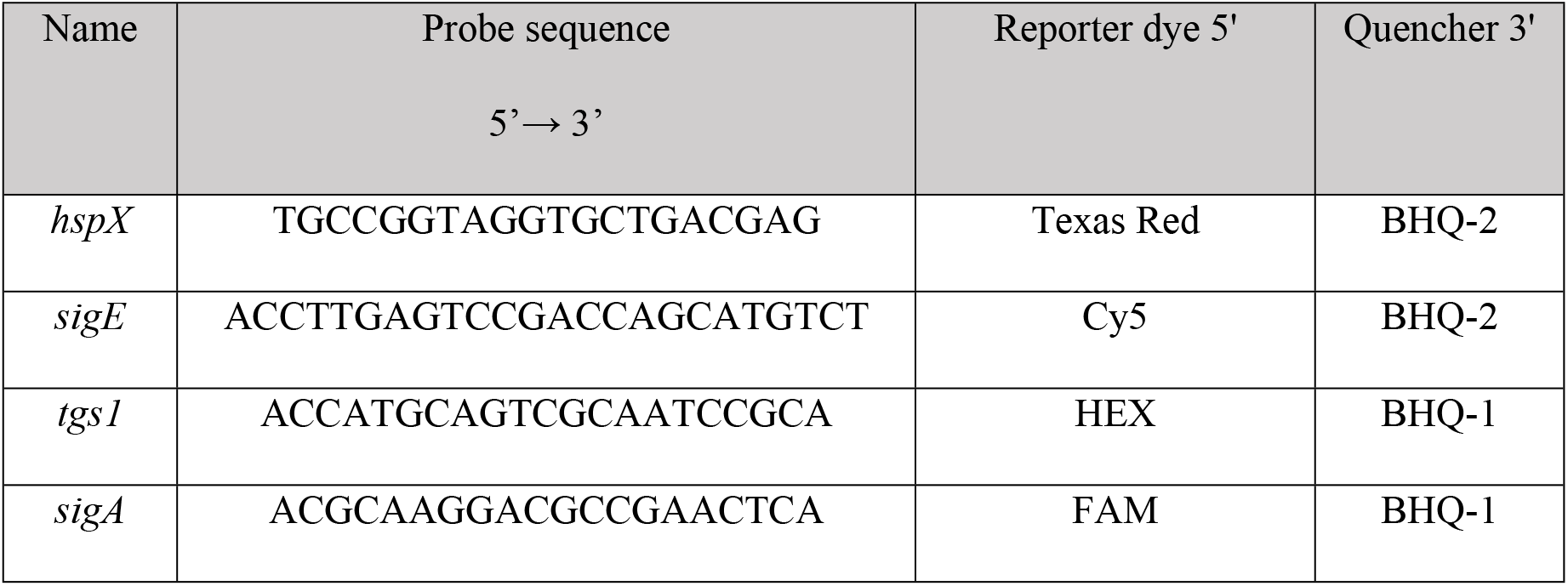
Probes sequences used in multiplex real-time PCR target stress-related genes.

**Table 6.** The condition for Multiplex real-time PCR with probe assay target stress-related genes.

| Steps | Temperature (°C) | Time | Cycles |
| --- | --- | --- | --- |
| Pre-denaturation | 95 | 1 min | 1 |
| Denaturation | 95 | 30 sec | 40 |
| Extension | 56 | 30 sec | 40 |

### Statistical analysis

The result was represented by the mean values of two or three independent experiments run in triplicate ± SEM (standard error means). Statistical analysis was performed in GraphPad Prism v8. One-way analysis of variance with Tukey’s multiple comparison was used to assess the data between different groups. Kruskal-Wallis was used when data are not normally distributed. *P*-value < 0.05 was considered statistically significant and * = *P* < 0.05, * * = *P* < 0.01, and * * * = *P* < 0.001.

## Results

### Genotypic and phenotypic drug resistance of MTB isolates

Drug-sensitivity test was conducted on MTB isolates using the GenoType MTBDRplus assay and agar proportion method to detect genotypic and phenotypic drug resistance profile of MTB, respectively. In the GenoType MTBDRplus assay, RIF resistance was detected using probes of the *rpoB* gene while INH resistance was detected using probes of the *katG* and *inhA* genes. The MUT1 band was observed in INH-R isolates that represent the mutation of the inhA regulatory region (C-15T). The MUT2B band was observed in RIF-R isolates and represents the mutation of rpoB at codon H526D. In MDR isolates, double mutations of katG and rpoB were found at codon S315T (MUT1 band) and D516V (MUT1 band), respectively (S1 Fig.). In the agar proportion method, mycobacteria were cultured with isoniazid, rifampin, streptomycin, ethambutol, and p-Nitrobenzoic acid. H37Rv was susceptible to all drugs, INH-R was resistant to isoniazid, RIF-R was resistant to rifampin, and MDR resistant to all drugs except p-Nitrobenzoic acid (S1 Table.). These results correlate with the GenoType MTBDRplus assay.

### The expression of ManLAM related genes was different in response to INH treatment

The specificity of primers was assessed using MTB isolates with different drug resistant profiles; H37Rv (Drug sensitive strain), INH-R, RIF-R and MDR. All primers tested produced a single band at a desirable product size, and no non-specific band was observed, demonstrating primer specificity (S1 File.). After INH treatment, the relative expression of ManLAM related genes was determined in all mycobacteria clinical isolates. Results were presented as relative expression, which is the fold-changes in ManLAM related gene expression normalized to *sigA* expression relative to untreated control.

The result of each ManLAM related gene was compared between four strains and presented in Fig 1. In H37Rv, only *mptA* that involved in elongation of LM was up-regulated (2.84) but the relative expression not significantly different among four strains (Fig 1B).

**Fig 1.**
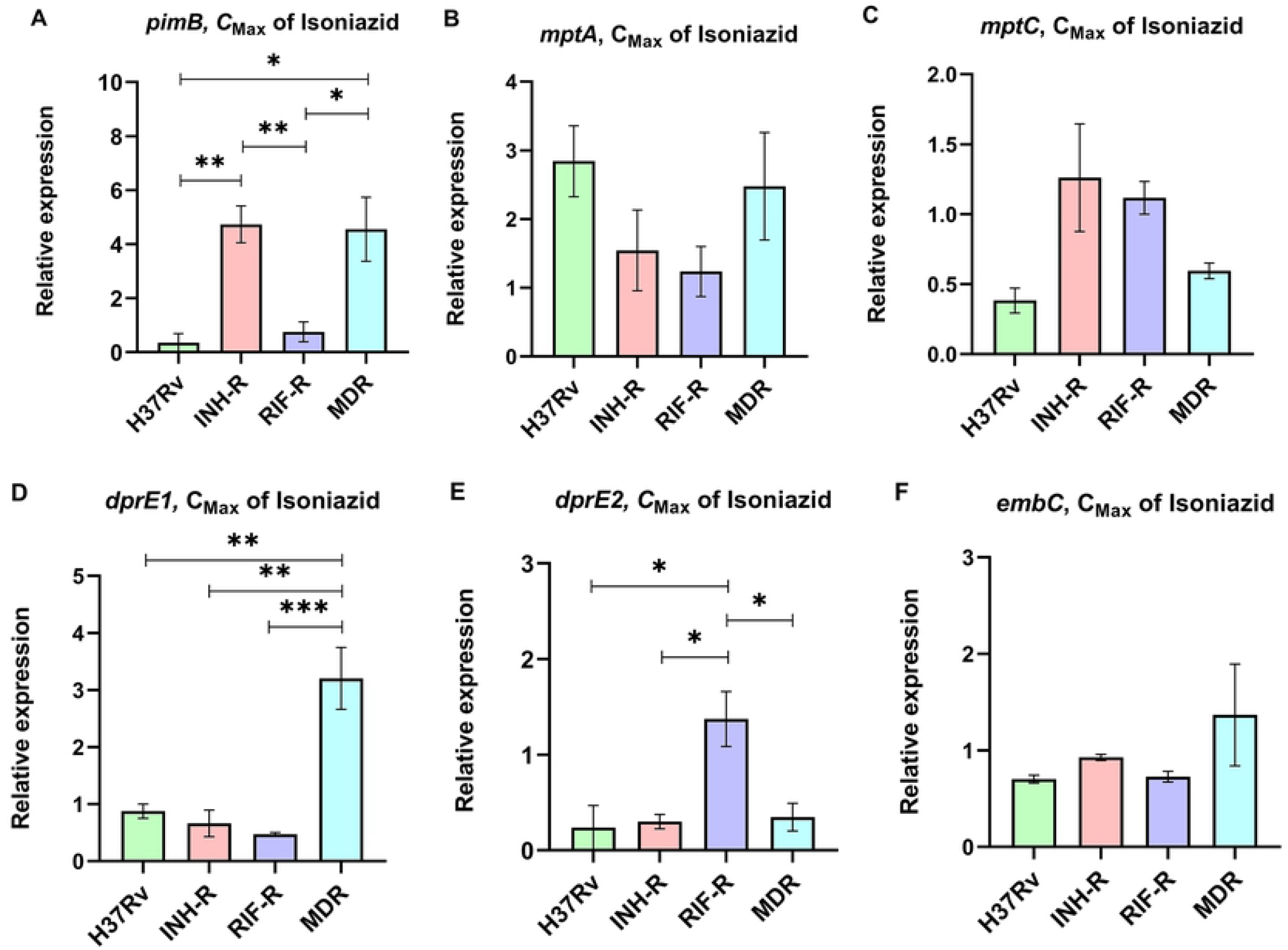
The relative expression of ManLAM related genes induced by INH treatment. The relative expression of (A) *pimB*, (B) *mptA*, (C) *mptC*, (D) *dprE1*, (E) *dprE2* and (F) *embC* of MTB fours strain following C_Max_ of INH treatment for 30 min.

Interestingly, in INH-R, *pimB, mptA*, and *mptC* were up-regulated. *pimB* is one of the genes responsible for PIMs production, which is involved in an early stage of ManLAM biosynthesis. PimB was up-regulated 4.73-fold and expressed significantly higher than H37Rv and RIF-R (*p*<0.01) (Fig 1A). *mptA* and *mptC* that are involved in elongation and branching of LM/LAM were up-regulated 1.54 and 1.26-fold, respectively (Fig 1B and 1C)

In RIF-R, ManLAM related genes were slightly up-regulated, including *mptA* (1.24-fold), *mptC* (1.12-fold), and *dprE2* (1.37-fold) (Fig 1B, 1C, and 1E).

In MDR, several genes were markedly up-regulated, including *pimB* (4.55-fold), *mptA* (2.48-fold) and *dprE1* (3.2-fold). MDR up-regulated *pimB* greater than H37Rv and RIF-R (*p*<0.05), while predominantly up-regulated *dprE1* among four strains. However, *embC* that involved in the biosynthesis of branched arabinan polymers on arabinogalactan was also upregulated in MDR by 1.37-fold, but not statistically significantly higher than others (Fig 1F).

ManLAM related genes that up-regulated was compared in each strain, only INH-R have a statistical significance result. *pimB* was strongly up-regulated greater than *mptA* and *mptC* in INH-R (Fig 2). Following experimental results, indicate that the expression pattern was significantly different between drug-sensitive (H37Rv) and drug resistant-TB strains (Fig 3). These results suggest that INH treatment induce ManLAM related genes expression, especially in drug resistant-TB strains.

**Fig 2.**
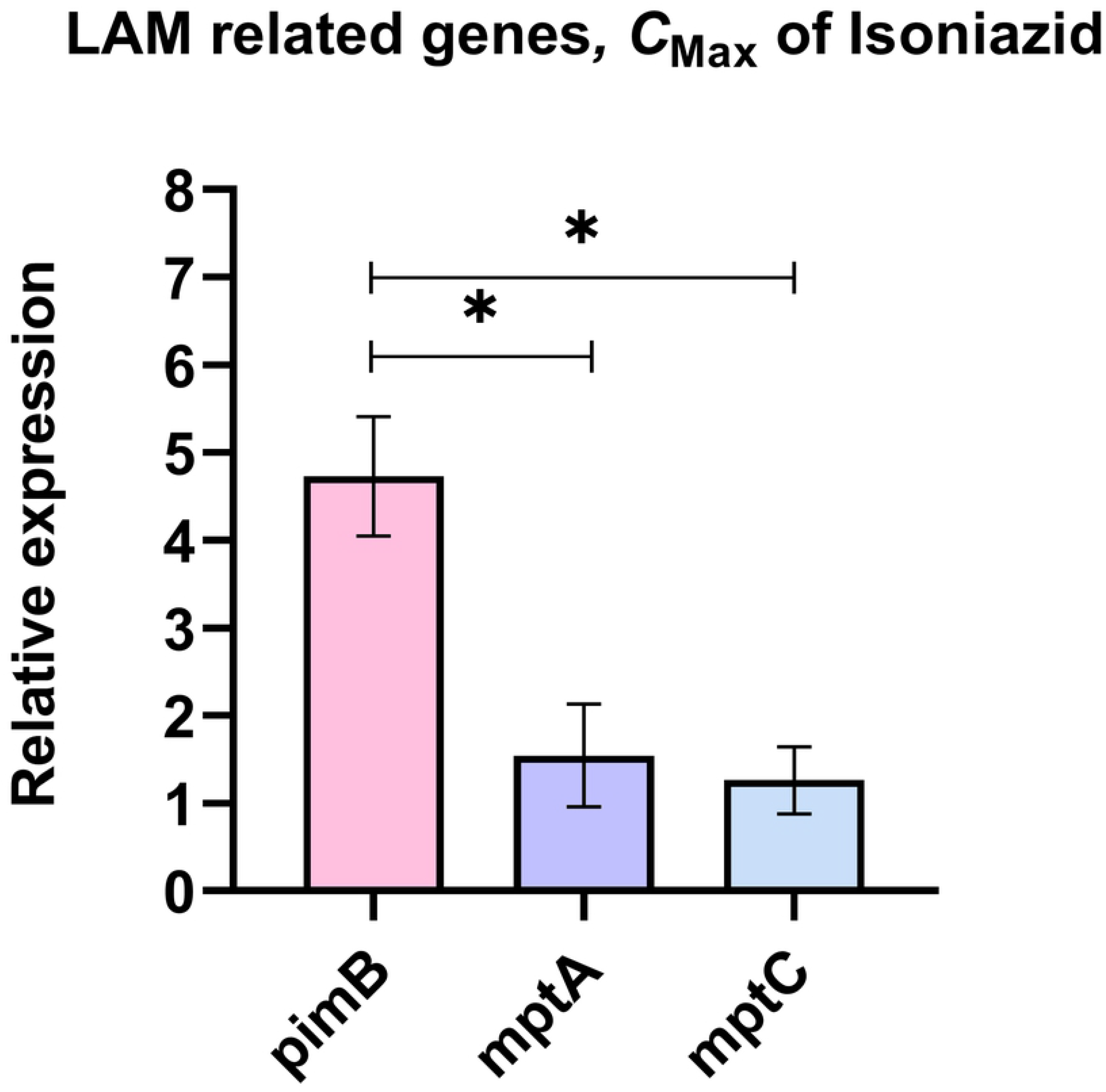
The relative expression of ManLAM related genes that were up-regulated in INH-R. The relative expression was compared among ManLAM related genes that were up-regulated in INH-R, including *pimB, mptA* and *mptC*.

**Fig 3.**
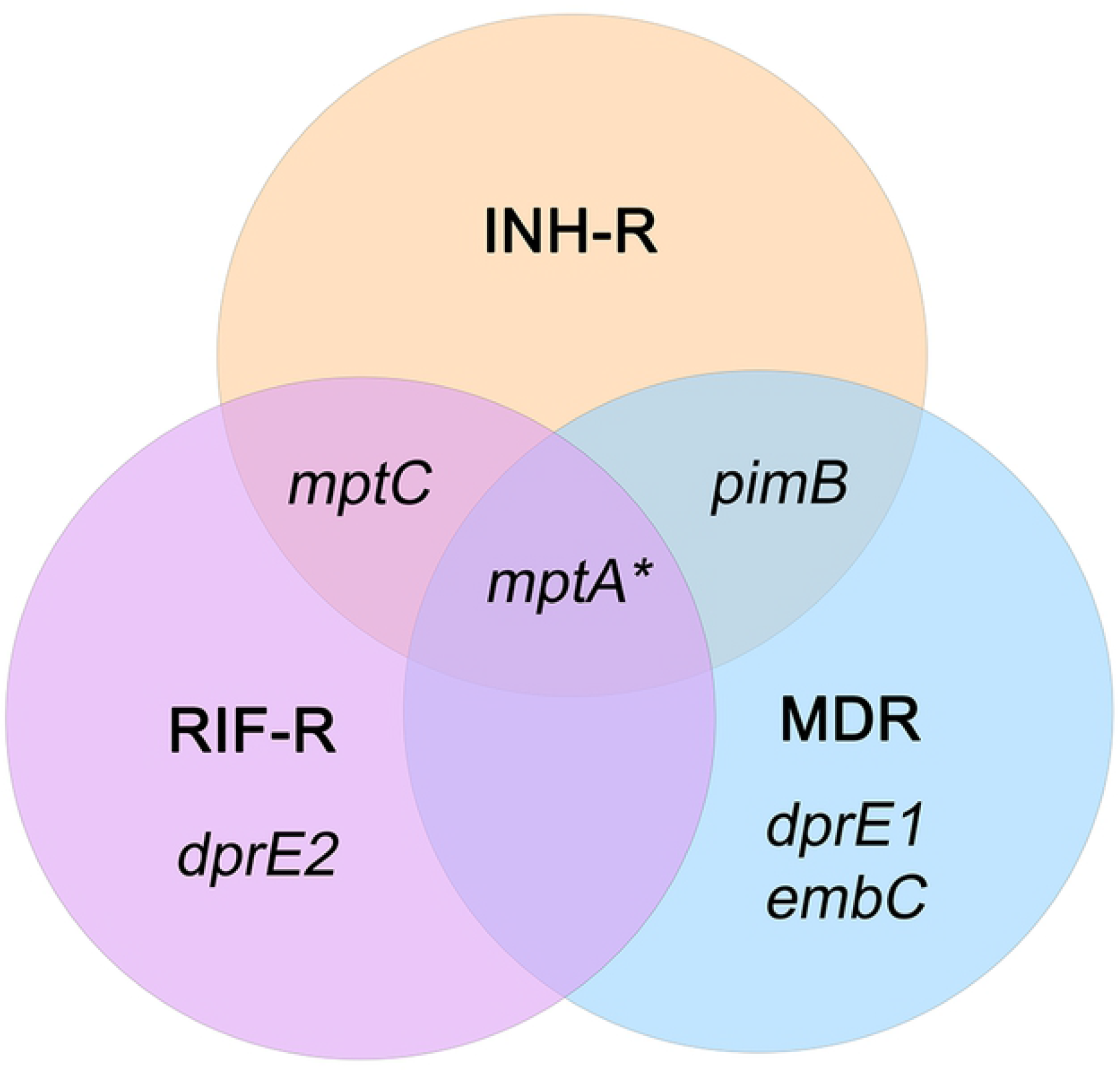
The unique expression pattern of ManLAM related genes of INH-treated MTB. ManLAM related genes that were up-regulated are listed in each circle that represents each MTB strain. An asterisk represents a gene which is up-regulated in H37Rv (a drug-sensitive strain).

### DNA sequencing of Drug resistance associated genes of MTB isolates

INH-R isolate showed a C-15T mutation at the *inhA* promoter region (Fig 4). The C-15T mutation causes *inhA* overexpression and requires higher doses of INH to complete inhibition [34]. In *rpoB* gene, a C→G mutation at nucleotide position 1333 was found in RIF-R, causing an amino acid substitution of aspartate (D) for histidine (H) at codon 445 (H526D, according to the *E. coli* numbering system) (Fig 5). MDR isolate showed a A→T mutation at nucleotide position 1304, causing an amino acid substitution of valine (V) for aspartate (D) at codon 435 (D516V, according to the *E. coli* numbering system) (Fig 5). Both *rpoB* mutations are mostly found in RIF-resistant strains and occur within an 81-bp region of the *rpoB* gene namely ‘RRDR’ (RIF resistance-determining region). Although the amino acid substitution of codon 526 confers high levels of RIF resistance, but mutation in codon 516 cause low levels of RIF resistance [35]. Moreover, MDR showed mutation of a G→C mutation at nucleotide position 944 of the *katG* gene, causing an amino acid substitution of Threonine (T) for serine (S) at codon 315 (S513T) (Fig 6)

**Fig. 4.**
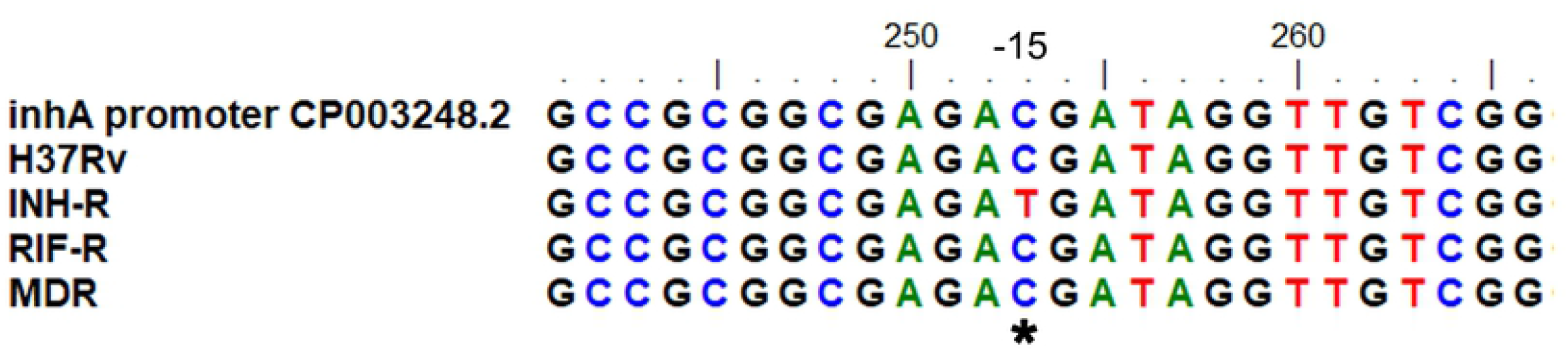
Multiple sequence alignment of *inhA* for MTB with different drug resistance profiles.

**Fig. 5.**
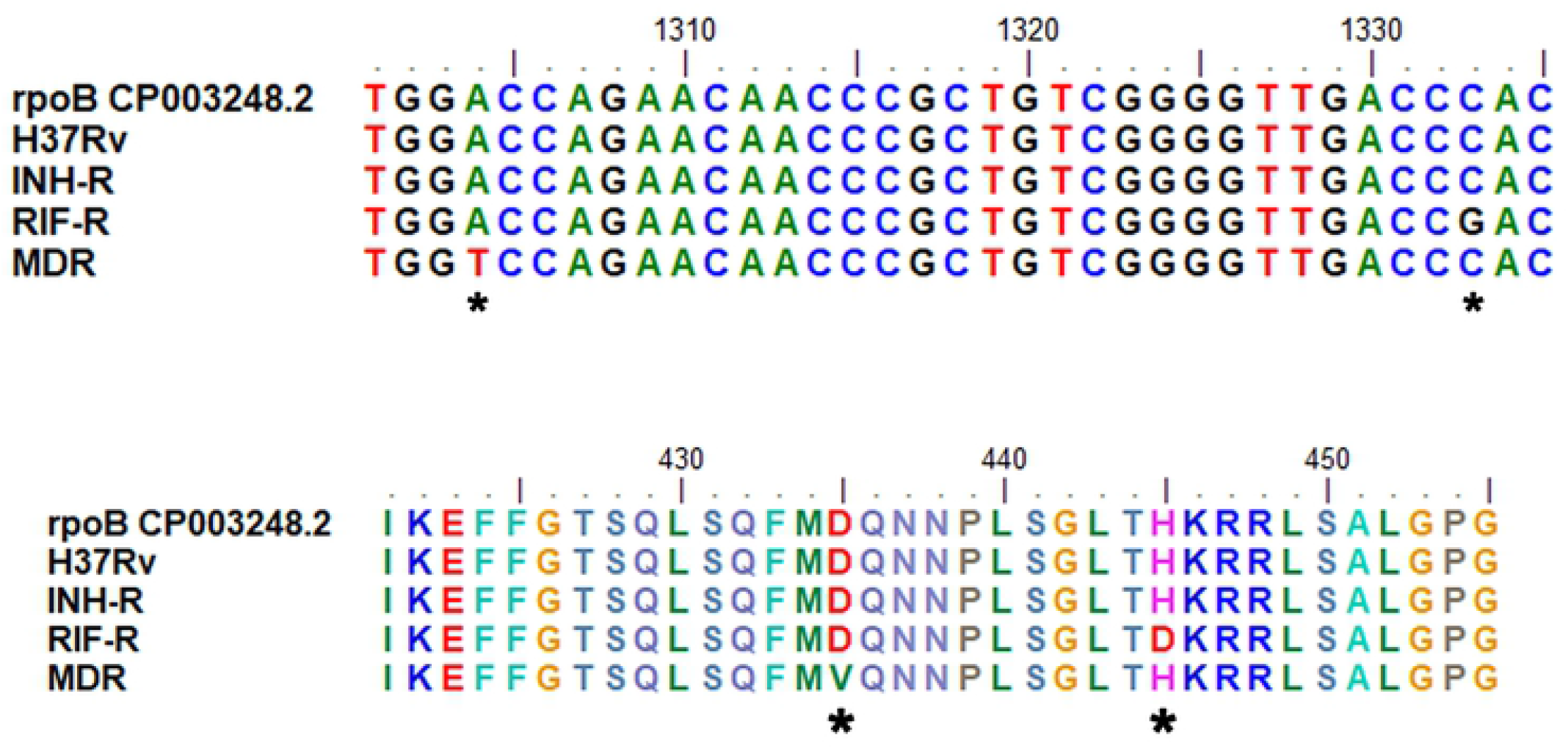
Multiple sequence alignment of *rpoB* for MTB with different drug resistance profiles.

**Fig. 6.**
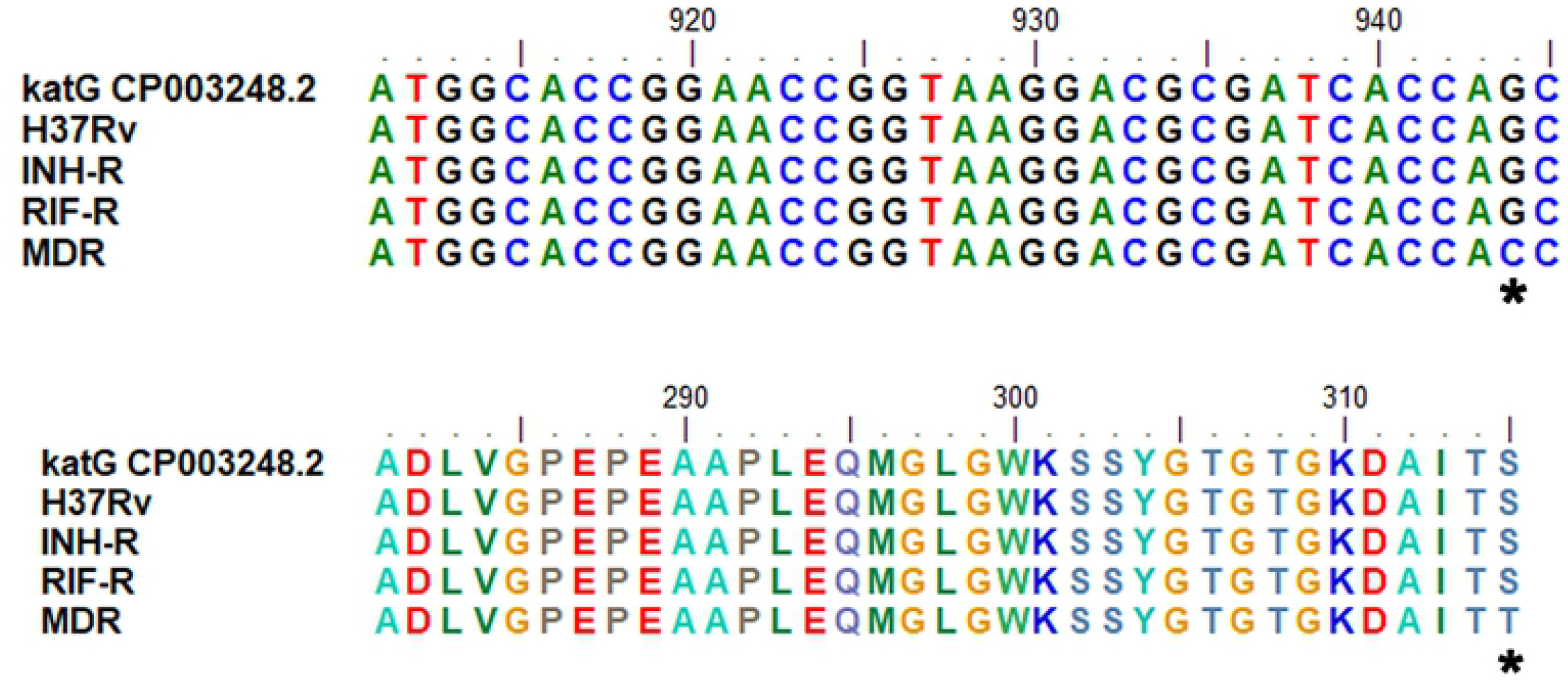
Multiple sequence alignment of *katG* for MTB with different drug resistance profiles.

### DNA sequencing of ManLAM related genes of MTB isolates

No differences in the DNA sequence of ManLAM related genes were observed, except for the *dprE1* gene. In RIF-R and MDR isolates that are resistant to rifampin showed *dprE1* mutation at nucleotide position 459 of the *dprE1* gene (C→T mutation), causing silent mutation or synonymous codon at position 153 (Fig 7). DprE1 is one of the TB drug targets, mutation of *dprE1* associated with MTB growth and drug resistance depend on type and position [36]. Codons that are translated to the same amino acid are called synonymous. Previous studies have reported that synonymous codons are not utilized in equal frequency over genes or genomes. This phenomenon, termed “codon usage bias” that may alter the expression level of the manipulated genes [37]. Codon usage for dprE1 of MTB four strains were analyzed by using visual gene developer 1.9 (Build 785) program. The frequency, number, and fraction of codons for each amino acid were evaluated. Almost codon usages in MTB four strains had similar frequencies, except ACC and ACT. In RIF-R and MDR, the fraction of ACT codon was higher compared to H37Rv while the ACC fraction was decreased (S2 File.).

**Fig. 7.**
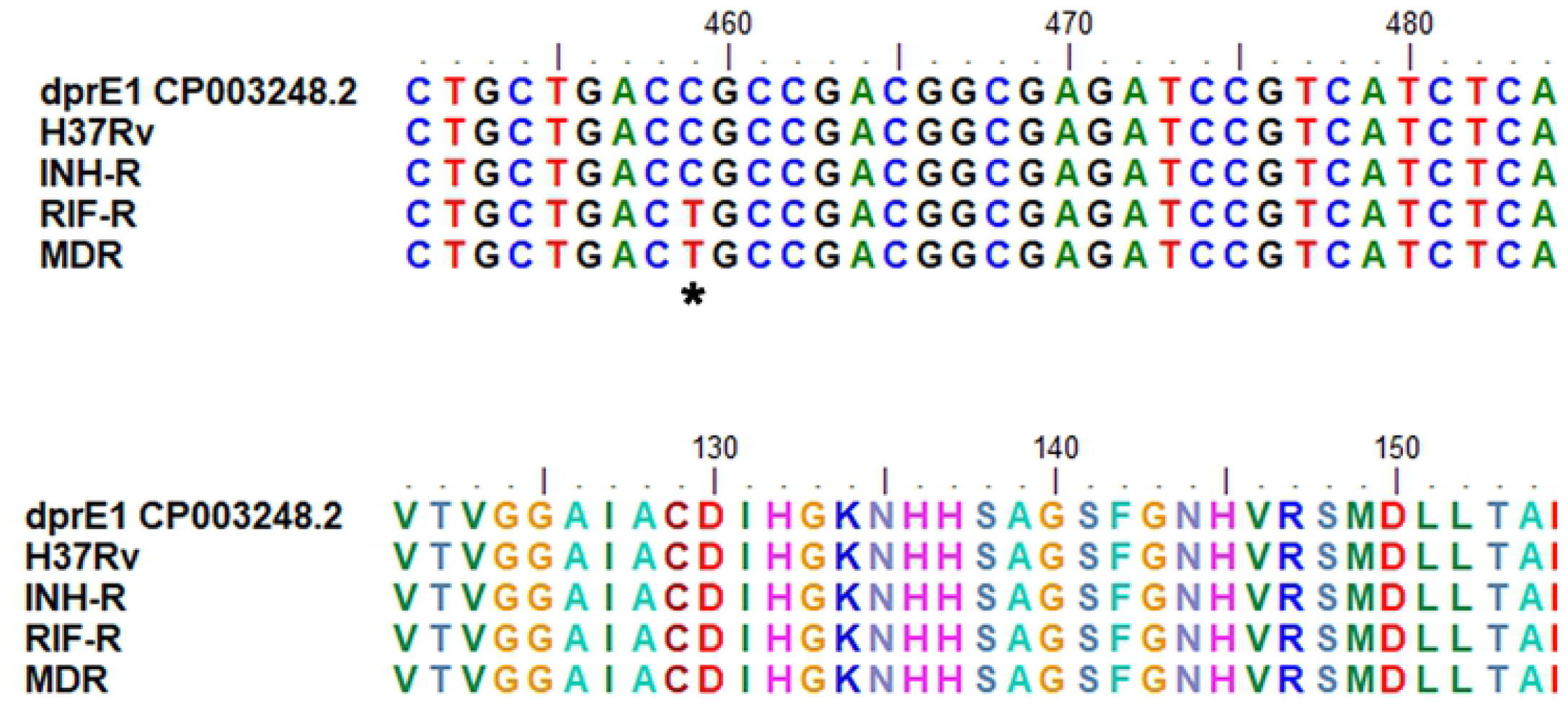
Multiple sequence alignment of *dprE1* for MTB with different drug resistance profiles.

### The expression of stress related genes was different in response to INH treatment and RIF treatment

Isoniazid (INH) and Rifampin (RIF) are first-line drugs for TB treatment. INH inhibits mycolic acid synthesis while RIF inhibits bacterial RNA polymerase. In this experiment, INH and RIF were used as stressors and *hspX, tgs1*, and *sigE* were used as stress responder markers. H37Rv, INH-R and MDR were treated with INH or RIF at C_Max_ concentration for 30 min to determine the effect of antibiotic stress. Results were presented as relative expression, which is the fold-changes in each stress-related gene expression normalized to *sigA* expression relative to untreated control. Primers and probes in this experiment were specific to the target gene and a non-specific band was not observed (S3 File.).

In response to isoniazid, MDR-MTB was significantly up-regulated *hspX* (1.42-fold) and *tgs1* (2.03-fold) greater than H37Rv and INH-R, while H37Rv (drug sensitive strain) was significantly up-regulated *sigE* (3.07-fold) greater than INH-R and MDR (*p*x< 0.01) as shown in Fig 8A-C.

**Fig 8.**
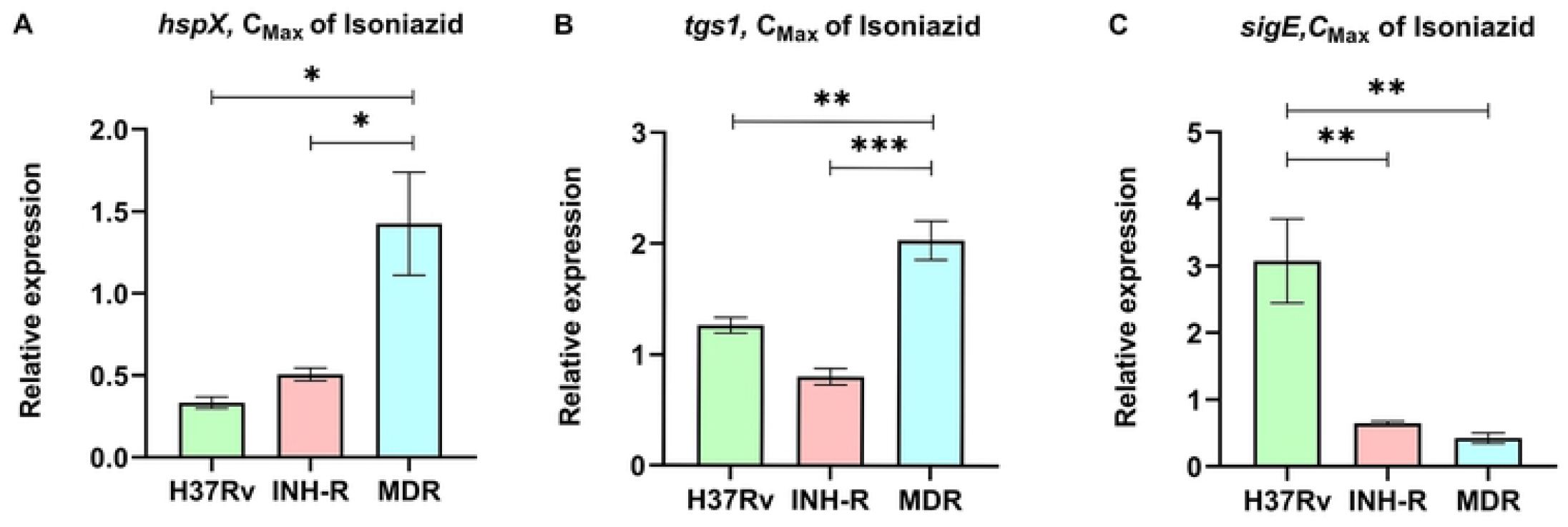
The relative expression of stress-related genes in INH-treated MTB. (A) *hspX*. (B) *tgs1*. (C) *sigE*.

In contrast to the results from INH treatment, *sigE* was significantly upregulated in INH-R (4.42-fold) and MDR (4.07-fold) in response to rifampin as presented in Fig 9A-C. However, the expression level of *hspX* and *tgs1* tended to increase in all strains but was not present at a statistically significant level.

**Fig. 9.**
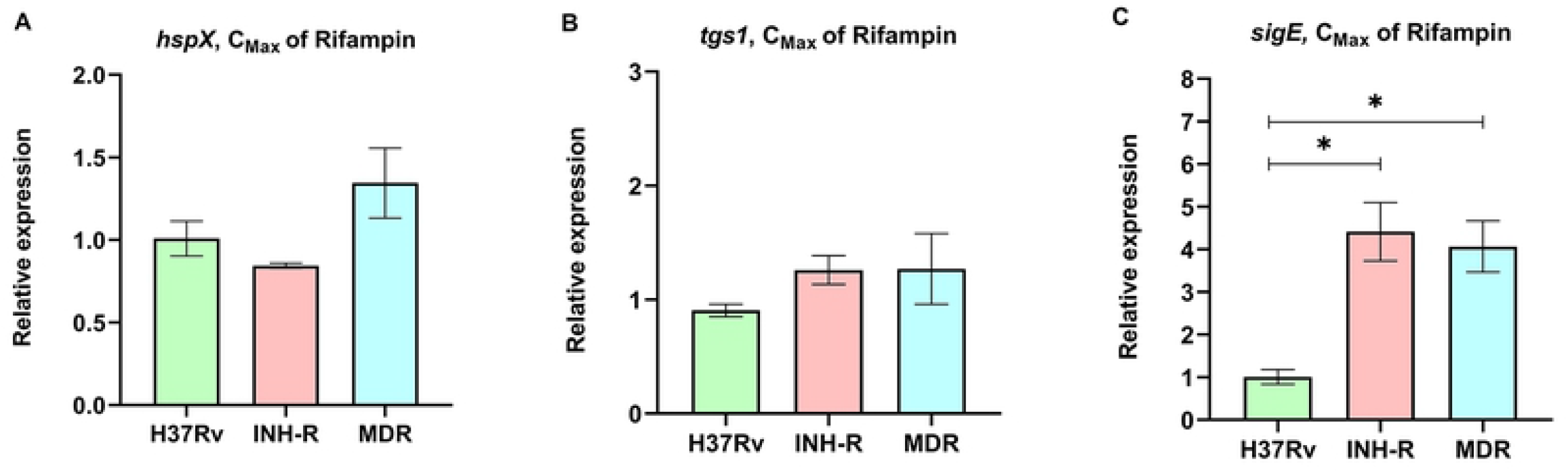
The relative expression of stress-related genes in RIF-treated MTB. (A) *hspX*. (B) *tgs1*. (C) *sigE*.

Previous publication categorized *hspX* and *tgs1* as dormancy regulon while *sigE* was classified as an enduring hypoxic response according to their function or expression pattern [19]. Both *hspX* and *tgs1* genes were clearly up-regulated in Wayne dormancy culture model (hypoxia). *sigE* was also up-regulated in this model but presented at a lower level compare to *hspX* and *tgs1* at the indicated time point. Similar result was observed, *hspX* and *tgs1* were significantly express in multi-stress model [12] [low oxygen, higher levels of carbon dioxide, nutrient starvation, and acidic pH (5.0)]. This multi-stress model induces MTB switch from an active to a dormancy phase, resulting in long-term survival and drug tolerance. Thus, these data suggest that environmental stress triggers MTB to regulate stress responder genes to promote their survival. A similar phenomenon may occur in the case of antibiotic stress such as INH or RIF, that shown in the up-regulation of *hspX, tgs1* and *sigE* in this work.

## Discussion

Previous publications reported that MTB responds to several stresses by remodeling cell wall components to promote their survival in different conditions, and these changes also affect their antibiotic susceptibility profile [12, 14, 18]. The model of cell wall regulation from a recent review article demonstrates that LM and LAM are moderately expressed in rapid growth but up-regulated during growth stasis [13]. Changes in LM and LAM structures significantly impact the cell wall integrity of *M. smegmatis*, resulting in loss of the acid-fast staining property, increased sensitivity to antibiotics, and more sensitive to macrophage killing. Similarly, defects in LM and LAM structures affect the pathogenesis of MTB [38]. Thus, these suggest that MTB regulated ManLAM expression, which is the virulence factor for protecting mycobacterial cells and supporting their long-term survival.

This study was interested in the stress response mechanism of *M. tuberculosis* induced by anti-TB drugs with a focus on INH. The aim of this study was to determine the expression level of ManLAM related genes and stress-related genes in response to INH in *M. tuberculosis* clinical isolates with various drug resistance profiles. Interestingly, ManLAM related genes tested in this study were regulated differently between each drug resistant strain, which causes a unique pattern. Almost all ManLAM related genes were up-regulated in drug-resistant strains, especially in MDR.

MDR isolates have katG (S513T) and rpoB (D516V) mutations, which are associated with INH resistance and RIF resistance, respectively. Even though INH-R and RIF-R have one phenotypic drug resistant profile like MDR, the pattern of LAM related gene expression was markedly different. INH-R has C-15T mutation in the inhA promoter region, while RIF-R has rpoB mutation (H526D). These results suggest that the genetic background is one of the factors involved in the adaptive response mechanism of MTB under INH stress.

Interestingly, INH-R and MDR were dramatically up-regulated *pimB. pimB* is one of the genes responsible for PIMs synthesis, the structural basis of ManLAM. PIMs play a role in maintaining cell wall integrity that provides mycobacteria with intrinsic antibiotic tolerance and modulates host response during infection [14]. PIMs activate cytokine production that induces granuloma formation and promotes disease progression [39, 40]. The expression level of mptA was up-regulated in all strains at similar levels, reflecting its requirement under INH stress conditions (Fig 1B). The balance activity of *mptA* and *mptC* is crucial for LM/LAM production in mycobacteria. Overexpression of MSMEG_4247, which is an orthologs of Rv2181(*mptC*) leads to the synthesis of dwarfed LM, shorter mannan backbone and smaller arabinan domain of LAM in *M. smegmatis* [38]. The expression level of *mptA* and *mptC* following INH treatment in this study was not significantly different in each strain except H37Rv (drug-sensitive strain), suggesting that the balance of these two genes’ expression also plays a role in the adaptive mechanism of MTB. The expression level of *dprE1* and *dprE2* was up-regulated in MDR and RIF-R, respectively. *dprE1* and *dprE2* are responsible for the epimerization of DPR to DPA [41], arabinose donor in the arabinan domains of arabinogalactan and ManLAM. DprE1 is essential in mycobacteria and it is a valuable drug target [42]. dprE1 depletion induces cell lysis, which results in cell death and is attenuated in macrophages [43]. DprE1 mutation affect mycobacterial growth and virulence, such as levels of resistance to drugs and changes in infectivity, depend on mutation type and position [36]. Alignment of RIF-R and MDR in this study showed a mutation at nucleotide position 459 of the *dprE1* gene that presents the same amino acid at codon 153. This synonymous codon used by RIF-R and MDR could be the result of codon usage bias, which can affect gene expression levels [37]. For dprE1, the ACT codon was preferred in RIF-R and MDR compared to the drug-sensitive strain. However, *dprE1* up-regulation was observed only in MDR, not RIF-R, suggesting that there are other factors that regulate *dprE1* expression response to INH treatment. *embC* (Rv3793) was up-regulated only in MDR. Previous work reported that *embC* is essential in MTB and the expression level of *embC* influenced the size of LAM in *M. smegmatis* but not in MTB. Moreover, the embC promoter was down-regulated under the hypoxia-induced nonreplicating condition, and *embC* was down-regulated in alveolar macrophages at an early stage of infection [10]. Thus, suggest that *embC* is regulated by mycobacteria depending on the type and duration of stressors.

The future study will determine the effect of INH stress on ManLAM under several stresses that will improve understanding of the stress response mechanism of mycobacteria.

## Abbreviations

INH-R: isoniazid-resistant strain
MDR: multidrug-resistant strain
RIF-R: rifampin-resistant strain
qRT-PCR: real-time quantitative reverse transcription polymerase chain reaction

## Acknowledgements

The authors would like to thank Faculty of Associated Medical Sciences, Chiang Mai University for their support with equipment.

## Supporting information

**S1 Fig. Genotypic drug resistance profiles of mycobacteria by GenoType MTBDRplus assay**. Lane 1; negative control, lane 2 and 3; MDR, lane 4 and 5; RIF-R, lane 6 and 7; INH-R, lane 8 and 9; H37Rv.

**S1 Table. Phenotypic drug resistance profiles of mycobacteria by Agar proportion method**. INH; isoniazid, RIF; rifampin, STM; streptomycin, ETM; ethambutol, PNB; p-Nitrobenzoic acid, S; susceptible.

**S1 File. Specificity of LAM related gene primers**. Fig A-C Electrophoresis of DNA amplified fragments obtained from the specificity test of MTB specific primers to targeted genes.

**S2 File. Codon usages for DprE1 of MTB with different drug resistant profile**.

**S3 File. Electrophoresis of PCR product using primers and probes target stress-related genes**.

## Notes

### Competing Interest Statement

The authors have declared no competing interest.

